# Secondary origin, hybridization and sexual reproduction in a diploid- tetraploid contact zone of the facultative apomictic orchid *Zygopetalum mackayi*

**DOI:** 10.1101/764134

**Authors:** Yohans Alves de Moura, Alessandro Alves-Pereira, Carla Cristina Silva, Lívia Moura de Souza, Anete Pereira de Souza, Samantha Koehler

**Affiliations:** Departamento de Biologia Vegetal, Instituto de Biologia, CP 6109, Universidade Estadual de Campinas – UNICAMP, 13083-970, Campinas, SP, Brazil

**Keywords:** apomixis, Brazil, hybrid, mixed-cytotype, reproductive interference, polyploidy, sympatry

## Abstract

>Mixed-cytotype populations are ideal to understand polyploid establishment and diversification. We used the orchid *Zygopetalum mackayi* to understand how facultative apomictic reproduction relates to polyploidy. Sexual diploids and facultative apomictic tetraploids occur under distinct niches, with a contact zone where triploids occur. We hypothesized that facultative apomictic reproduction increases the fitness of tetraploids through reproductive interference between cytotypes. We predict patterns of genetic diversity of allopatric tetraploid populations to be significantly different from contact zone populations as a result of dominant apomictic reproduction in the later. We also describe the contact nature of diploids and tetraploids and the role of the intermediate triploids based on patterns of genetic structure within and among pure and mixed-cytotype populations.
>We designed eight microsatellite markers and genotyped 155 individuals from six populations resulting in 237 alleles. We described patterns of genetic diversity and structure within and among populations and cytotypes.
>Genotypic diversity is similarly high among all populations and cytotypes. Each cytotype emerged as a genetically cluster, combining individuals from different populations. Triploids clustered in an intermediate position between diploids andtetraploids.
>We rejected the hypothesis of reproductive interference between cytotypes of *Z. mackayi*. Patterns of genetic diversity are incongruent with the occurrence of apomict reproduction in tetraploids. Mixed-cytotype populations originate from secondary contact and triploids are hybrids between diploids and tetraploids and act as a reproductive barrier. We suggest polyploidy rather than facultative apomixis explains higher fitness of tetraploids in this species and, therefore, eco-geographical patterns of distribution.

## INTRODUCTION

Populations composed by two or more cytotypes (i.e. individuals with different numbers of chromosomes in their somatic cells) are ideal to understand the role of polyploidy in lineage diversification (Kolář *et al.* 2017). The physical proximity of different cytotypes allow inference on the magnitude of inter-cytotype reproductive isolation and of conditions for polyploid establishment (e.g. Trávnícek *et al.* 2011; Kolář *et al.* 2012). Therefore, describing the patterns of genetic diversity within and among cytotypes is a valuable tool to understand the origin of polyploid populations, the competitive dynamics among cytotypes, and the effect of distinct reproductive strategies on partitioning genetic diversity (Karunarathne *et al.* 2018).

Mixed cytotype contact zones can be formed when polyploids arise and coexist with their direct progenitors in a primary contact zone. Alternatively, distinct cytotypes can evolve in allopatry and range expand to a secondary contact zone (Petit *et al.* 1999; Husband *et al.* 2013). When triploids occur in primary or secondary mixed-cytotype populations they usually act as a barrier to gene flow between populations with distinct ploidies because of their reduced adaptability (Ramsey & Schemske 1998) or because of a triploid block (i.e. massive abortion of triploid seeds caused by failure of the endosperm tissue, Marks 1966; Köhler *et al.* 2010). The production of low fitnesstriploids is expected to cause the exclusion of rare and unfit new cytotypes in a frequency-dependent process called minority cytotype exclusion (MCE) (Levin 1975; Felber 1991, Husband 2004). However, when fertile or partially fertile triploids exist, they may promote coexistence of multiple cytotypes through a triploid bridge (i.e. union between haploid gametes produced by diploids and unreduced triploid gametes produced by triploids, resulting in tetraploids, Ramsey & Schemske 1998), even when they are rare (i.e. Verduijn *et al.* 2004).

Asexual reproduction is usually correlated with polyploidy and within-population cytotype diversity (Kolář *et al.* 2017). Reproductive interference results from negative effects of any interspecific sexual interaction on fitness (Ribeiro & Spielman 1998), such as intercytotype crosses between sexual diploids and higher-ploidy apomicts. Apomictic cytotypes capture pollen from exclusively sexual cytotypes, decreasing its reproductive success, whereas intermediate cytotype progeny produced with diploid pollen compete with polyploid apomictic embryos and, therefore, rarely succeed (Hersh *et al.* 2016). Previous attempts to identify whether intrinsic effects related to changes in the reproductive mode or in ploidy drive eco-geographical differentiation yield various results. Mau et al. (2015) concluded that ploidy, rather than mating system, determines a shift in ecological niches in *Boechera* spp., while Alonso-Marcos et al. (2019) found the opposite for *Potentilla puberula* and Schinkel et al. (2016) found no difference in ecological conditions of co-ocurring apomictic and sexual individuals of *Ranunculus kuepferi*. Ecological diversification of sexuals and apomicts were mainly explored in previously glaciated areas of the northern hemisphere (Hojsgaard & Hörandl 2015). The transition from sex to apomixis is frequently associated with a major change in geographical distribution (i.e. geographical parthenogenesis) in which apomictic lineages occupied new environments that were severely affected by the glacial cycles of the Late Pleistocene (Kearney 2005). In the tropics, the impact of glaciations on diversification patterns was far more complex, involving changes in elevation ranges and expansion-retraction dynamics (Markgraf *et al.* 1995).

The orchid *Zygopetalum mackayi* Hook. is a good model to understand how facultative apomictic reproduction relates to polyploid ecological diversification in the tropics. Diploid (2N = 48) and tetraploid cytotypes (2N = 96) of *Z. mackayi* have a parapatric distribution and occur under different climatic conditions, meeting in a contact zone where triploids (2N = 72) are found (Gomes *et al.* 2018). Tetraploids have a much larger distribution than diploids (Gomes *et al.* 2018). Populations dominated by tetraploids occur in areas of high annual precipitation and marked temperature, whereas populations dominated by diploids are in areas of low annual precipitation and subtle temperature seasonality. Mixed-ploidy populations occur in areas of intermediate climatic conditions (Gomes *et al.* 2018). Triploids and tetraploids are facultative apomictic, producing sexual and apomictic embryos in the same fruit (Afzelius 1959). Apomixis does not provide reproductive assurance in *Z. mackayi*. Although this species is self-compatible, it is dependent on large bees for fruit formation (Campacci *et al.* 2017).

In this study, we test the hypothesis that facultative apomictic reproduction in *Z. mackayi* increases the fitness of tetraploids through reproductive interference. We predict patterns of genetic diversity of pure tetraploid populations to be significantly different of mixed-cytotype populations as a result of dominant apomictic reproduction in the contact zone. We also describe the contact nature of diploids and tetraploids and the role of the intermediate triploids based on patterns of genetic structure within and among pure and mixed cytotype populations of the orchid *Z. mackayi*.

## MATERIAL AND METHODS

### Sampling and DNA extraction

One hundred and fifty-five individuals were sampled from six populations in and around the contact zone of cytotypes based on the cytogeographic study of Gomes *et al*. (2018). Populations NB-2X and PD-2X are represented only by diploids, populations GAR-4X and MAR-4X only by tetraploids and populations ANG-M and PI-M by all three cytotypes, including triploids (Table 1, Fig. 1). Cytotypes were sampled for leaf tissue, identified and vouchered in the study of Gomes et al. (2018). Total DNA was extracted with the NucleoSpin® Plant II kit (Macherey-Nagel, Düren, Germany) or the DNeasy Plant Mini Kit (QIAGEN Biotecnologia Brasil Ltda, São Paulo, Brazil).

**Table 1.**
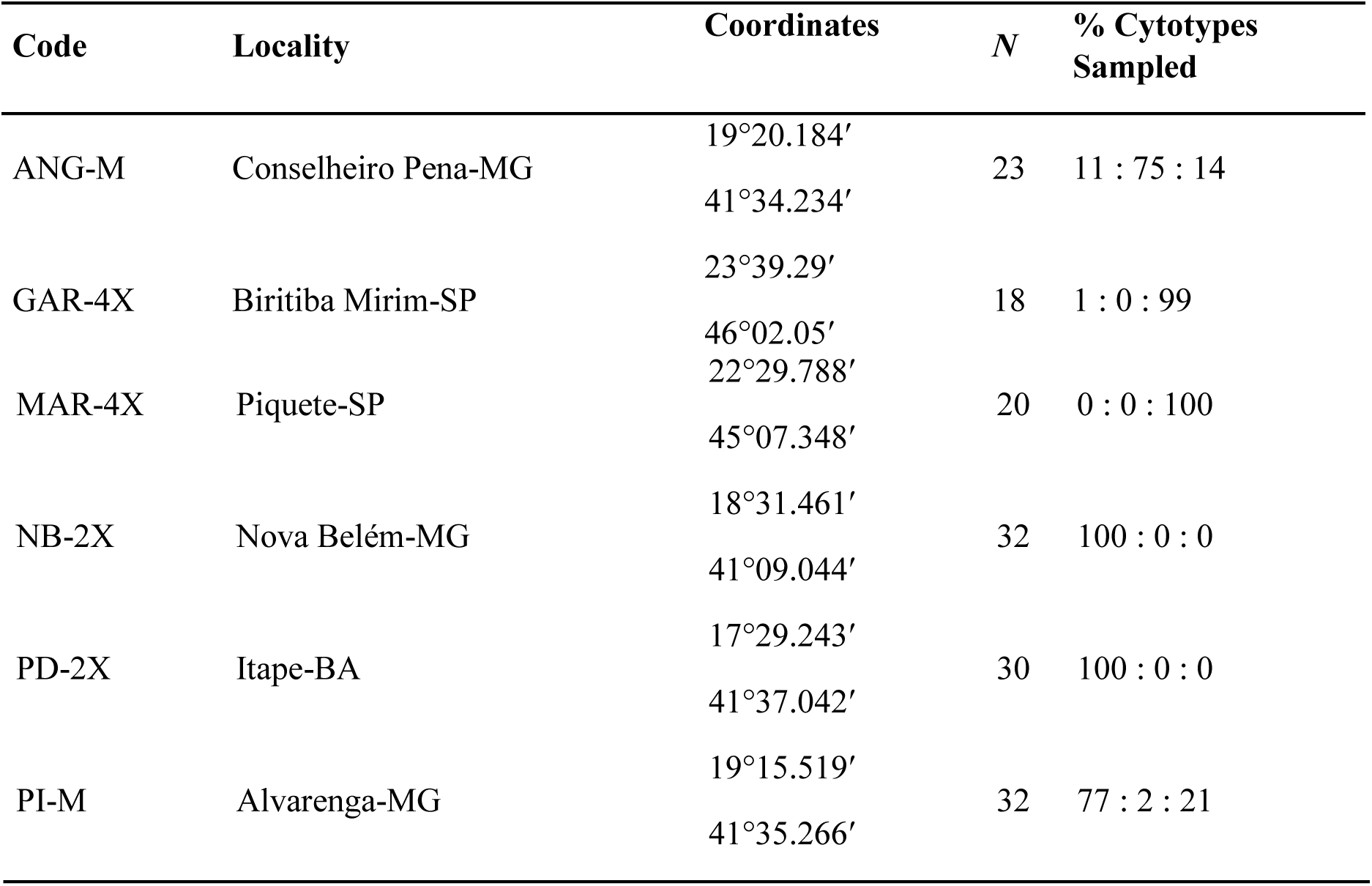
Sampling site code, locality, sample size (*N*), percent of diploids, triploids, and tetraploids of *Zygopetalum mackayi* Hook. sampled in each locality.

**Figure 1.**
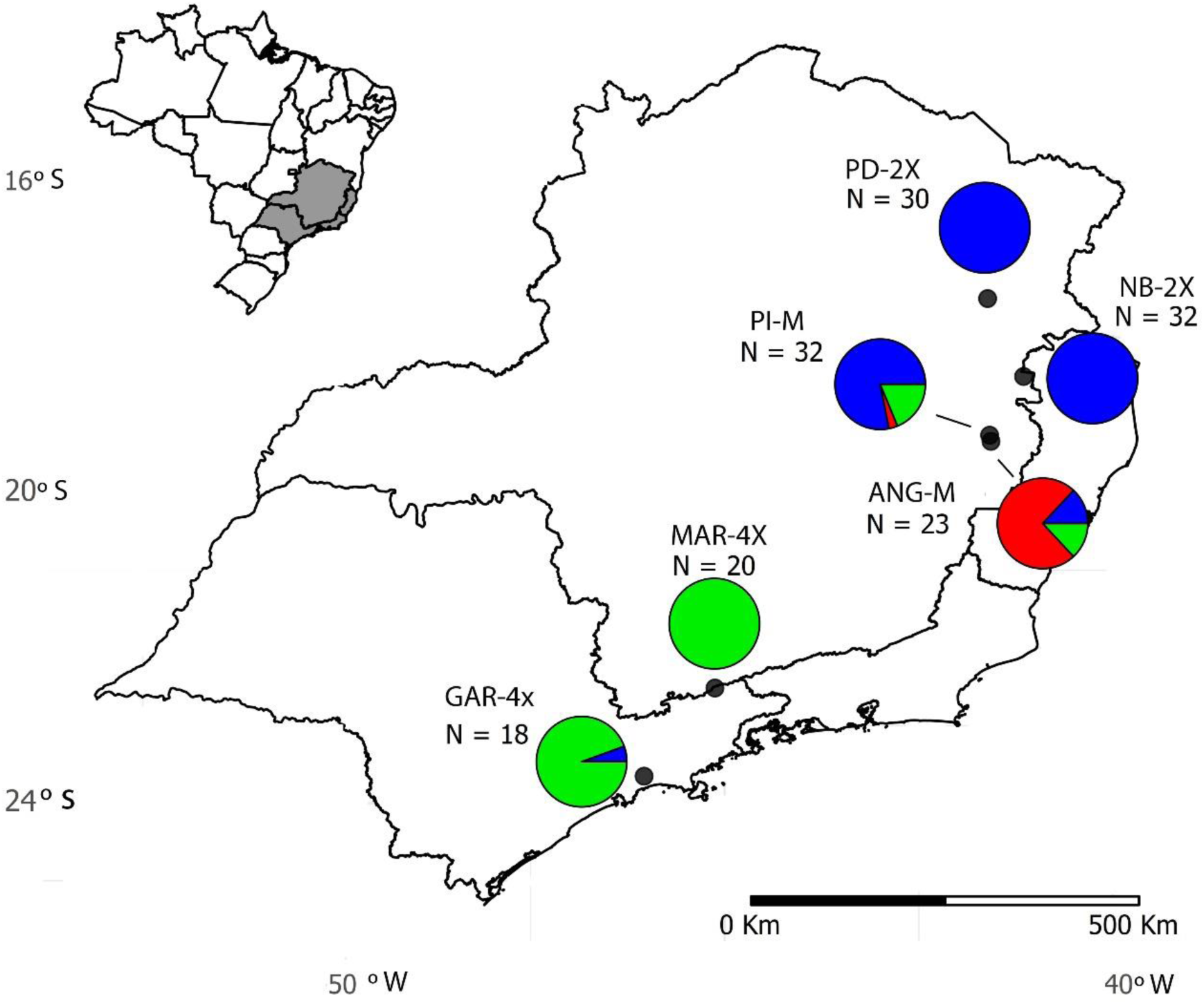
Geographic location of populations of *Zygopetalum mackayi* Hook. used in this study with cytotypes sampled per population represented by pie charts. The proportions of cytotypes in each population are represented by different colors (blue = diploid, red = triploid, green = tetraploid), (N) = number of individuals sampled.

### Microsatellite library development

Six microsatellite-enriched libraries for each sampled population of *Z. mackayi* were prepared following Billotte *et al.* (1999) with modifications. Libraries were based on two diploid specimens from populations NB-2X and PD-2X, one triplod specimen from population ANG-M and three tetraploid specimens from populations GAR-4X, MAR-4X and PI-M. Voucher specimens for each library were deposited at the herbarium UEC (Universidade Estadual de Campinas, Brazil). Genomic DNA was digested with the enzyme *Rsa1* (Integrated DNA Technology-IDT, Coralville, USA) and the fragments resulting from digestion were linked to *Rsa21* and *Rsa25* adapters. Fragment-containing repeats were selected with (CT)8 and (GT)8 biotinylated oligos and recovered with streptavidin-coated magnetic particles (Streptavidin MagneSphere Paramagnetic Particles, Promega, Madison, USA). Enriched DNA fragments were amplified and cloned using the pGEM-T easy vector (Promega, Madison, USA) and transformed into XL1-BLUE *Escherichia coli* competent cells (Stratagene, Santa Clara, USA). Three hundred and thirty-six clones (48 from each library) were sequenced using universal T7 and SP6 primers with a BigDye v3.1 terminator kit on an ABI 3500XL Genetic Analyzer automated sequencer (Applied Biosystems, Foster City, USA).

The sequences had the vectors removed in the software Chromas (http://www.technelysium.com.au/), submitted to quality selection and conversion to the format FASTA in cap3 (Huang & Madan 19xs99). The microsatellite motifs were searched using Gramene ssr finder tool (Temnykh *et al*. 2001). Primers were designed in primer3plus (Untergrasser *et al.* 2007) according to the following parameters: 18 - 26 bp primers; melting temperature (Tm) between 45 - 65 oC, with up to 3 oC of difference between each oligo in a pair of primers; salt concentration of 50 mM; GC content between 40 - 60%; fragment length between 100 - 360 bp. (Table S1)

Eight pairs of microsatellite primers were selected after a pilot study considering all 56 designed primers and one individual from each of the six populations (Table S2). The polymerase chain reaction (PCR) consisted of an initial denaturation at 95 °C for 5 min; 10 cycles of 95 °C for 1 min; with a touchdown from 63 °C to 54 °C (−0.5 °C/ cycle) for 1 min; 72 °C for 1 min; 25 cycles of 95 °C for 1 min; 52 °C for 30 s; 72 °C for 1 min; and a final extension at 72 °C for 5 min. The amplification mixture contained 20 ng of total DNA, 50 mM KCl, 20 mM Tris-HCl (pH 8.4), 1.5 mM MgCl2, 2 mM dNTPs, 1 mM of each primer (forward and reverse) and 1U of *Taq* polymerase. The success of amplifications of SSR loci was checked on agarose gel (2-3% w/v). For genotyping, PCR products were separated on 6% denaturing polyacrylamide gels for 3 hours at 250 volts and visualized by silver staining (Creste *et al*. 2001). For each locus, polymorphism information content (PIC) (Botstein *et al.* 1980) was calculated using the *polysat* (Clark & Schreier 2017) for R 3.4.0 (R Development Core Team 2017).

### Genetic and genotypic diversity

Analyses were first performed transforming the multilocus data into binary arrays of168 presence or absence of each allele for each individual. This data coding approach is169 commonly employed to genetic diversity studies in polyploids in which patterns of genetic inheritance are unknown (e.g. Sampson & Byrne 2011; Servick *et al*. 2015). The genetic parameters estimated for each population were: total number of alleles (*Na*),172 number of private alleles (*Np*) and average ratio of heterozygotes per locus (*Hp*).173 Complementary, the number of multilocus genotypes (*MGL*) was calculated to174 approximate clonal (apomictic) data to panmictic populations and to remove the effect of genetic linkage (Kamvar *et al*. 2014). This approach combines all alleles at each locus for each individual as a unique genotype to calculate the number of observed genotypes (richness), diversity and evenness. Genetic diversity within and among populations were estimated from three distinct indices as they accommodate the type of data used (allelic diversity): Shannon index (Shannon 2001), Simpson index (Simpson 1949) and Nei index (Nei 1978). Parameter values were estimated using the *poppr* and181 *polysat* packages (Kamvar *et al.* 2014; Clark & Schreier 2017) for R 3.4.0 (R Development Core Team 2017).

Analyses considering only diploid cytotypes and codominant, non-clonal data were also performed. Genetic diversity parameters of observed (*Ho*) and expected heterozygosity (*He*) and the inbreeding coefficient (*f*) were estimated using the *diversity* package (Keenan *et al*. 2013) for R 3.4.0 (R Development Core Team 2017). Recent population bottleneck signs were also evaluated for diploid cytotypes using BOTTLENECK 1.2.02 (Piry *et al*. 1999), considering deviations from the mutation-drift equilibrium. Two tests were used considering the indication of bottlenecks in the presence of significant excess heterozygosity: “Sign test” (Cornuet & Luikart 1996) and the “Wilcoxon sign-rank test” (Luikart & Cornuet, 1998), both based on the Infinite Alleles Model (IAM) and Stepwise Mutation Model (SMM) (10.000 iterations, P-value < 0.05).

### Genetic structure

The genetic relatedness among cytotypes was estimated based on Bruvo distances, using the package *polysat* (Clark & Schreier 2017) for R 3.4.0 (R Development Core Team 2017). Bruvo distances allow the comparison of individuals with different ploidy levels (Bruvo *et al*. 2004). The nature of the contact between cytotypes (primary vs. secondary) was then evaluated using a discriminant analysis of principal components (DAPC), which does not rely on specific genetic models to infer genetic structure, hence, well suited for data which include individuals of different ploidy levels (Jombart *et al*. 2010). Analyses were performed in using the *find.clusters* function of *adegenet* package (Jombart *et al.* 2008) for R 3.4.0 (R Development Core Team 2017). Clusters were estimated first considering all individuals together and then considering only individuals within each cytotype.

Two contrasting models explaining genetic relationships among cytotypes andpopulations were also evaluated with an analysis of molecular variance (AMOVA) (Excoffier *et al.* 1992) using the *poppr* package (Kamvar *et al.* 2014) for R 3.4.0 (R Development Core Team 2017). In a multiple origin model (A), each cytotype originates independently and repeatedly in each population. This hypothesis specifies a hierarchical AMOVA model in which genotypes are nested within cytotypes and cytotypes within populations. Alternatively, in a single origin model (B), each cytotype originates once and represent a distinct genetic group. In this model, genotypes were nested within populations nested within cytotypes. The favored AMOVA model is that in which the higher level (cytotypes for the single origin model or populations for the multiple origins model) explains the greatest component of total genetic variation. The significance of variance components was tested with 19999 permutations. We applied the information criterion of Akaike for model selection (AICc; Burnham & Anderson 1998).

Pairwise estimates of *Gst* (Nei & Chesser 1983) and their significance based on 10000 permutations were calculated to evaluate population differentiation using the *polysat* package (Kamvar *et al*. 2014). The estimates were plotted in a *heatmap* using the package *heatmap3* (Zhao *et al.* 2014) for R 3.4.0 (R Development Core Team 2017).

## RESULTS

### Genetic and genotypic diversity

In total, 237 alleles were amplified from the eight microsatellite loci, 114 in diploids, 38 in triploids, and 85 in tetraploids. The number of alleles amplified per locus ranged from 16 at locus GARD2 to 40 at locus PI-1 (Table S3). Diploids and tetraploids shared 34 alleles (14%), diploids and triploids shared 24 alleles (10%) and triploids and tetraploids shared 20 alleles (8%). We identified 20 private alleles (8.4 %) in diploids, 9 in triploids (3.8%) and 12 in tetraploids (5.1%).

Estimates of genetic diversity for the analyzed populations are listed in Table 2. Intrapopulation genetic diversity (N*a*, N*p*, H*p, MLG*, diversity indexes) was similar for the mixed cytotype populations (ANG-M and PI-M) in comparison to single cytotype populations (GAR-4X, MAR-4X, NB-2X and PD-2X) (Table 2). Most individuals had distinct multilocus genotypes within and among populations. Only two individuals from population GAR-4X shared the same MLG (Table 2).

**Table 2.** Estimates of within-population genetic diversity obtained with dominant data of eight nuclear microsatellites in *Zygopetalum mackayi* Hook.: population codes are according to Table 1, number of different alleles (*Na*), number of private alleles (*Np*), proportion of observed heterozygotes and standard deviation (*Hp*), number of multilocus genotypes (*MLG*), expected MLG considering the smallest population sampled in this study (*eMLG*), Shannon diversity index (Shannon 2001); Simpson diversity index (Simpson 1949), Nei diversity index (Nei 1978).

| Code | N | Na | Np | Hp | MLG | eMLG | Shannon index | Simpson index | Nei index |
| --- | --- | --- | --- | --- | --- | --- | --- | --- | --- |
| ANG-M | 23 | 44 | 9 | 0.576+/-0.24 | 23 | 18 | 3.140 | 0.969 | 0.188 |
| GAR-4X | 18 | 42 | 7 | 0.868+/-0.15 | 17 | 17 | 2.810 | 0.938 | 0.303 |
| MAR-4X | 20 | 36 | 5 | 0.592+/-0.25 | 20 | 18 | 3.000 | 0.950 | 0.113 |
| NB-2X | 32 | 38 | 8 | 0.617+/-0.28 | 32 | 18 | 3.470 | 0.967 | 0.144 |
| PD-2X | 30 | 30 | 7 | 0.425+/-0.20 | 30 | 18 | 3.400 | 0.957 | 0.296 |
| PI-M | 32 | 47 | 5 | 0.636+/-0.20 | 32 | 18 | 3.470 | 0.969 | 0.277 |
| Global |  |  |  |  | 154 |  | 5.030 | 0.993 | 0.293 |

Analyses of genetic diversity considering only diploids reveled high genetic diversity, no evidence of inbreeding and an excess of heterozygotes for the three analyzed populations (NB-2X, PD-2X and PI-M) (Table 3). Analysis with the software BOTTLENECK did not show evidence for a recent bottleneck in any of the diploid populations (NB-2X, PD-2X and PI-M) (P > 0.05 for all test results) (Table S4).

**Table 3.**
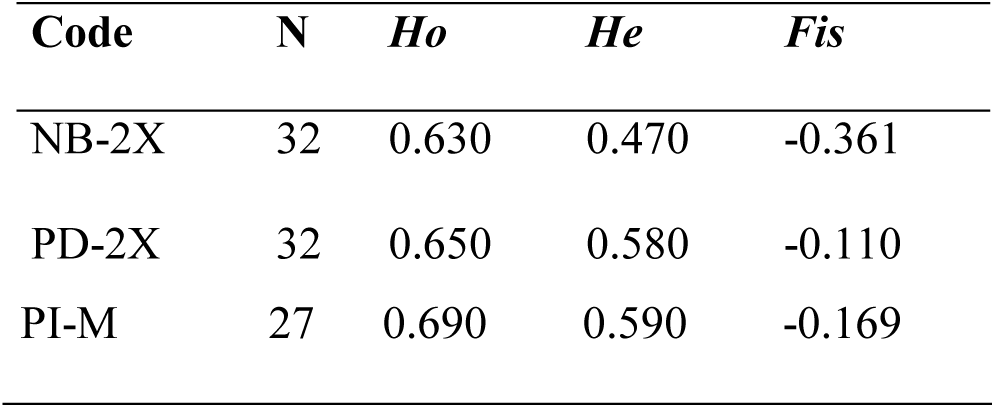
Estimates of within-population genetic diversity and inbreeding based on codominant data of eight nuclear microsatellites in diploids of *Zygopetalum mackayi* Hook. Population codes are according to Table 1, sample size (*N*), observed heterozygosity (*Ho*), expected heterozygosity (*He*), inbreeding coefficient (*Fis*).

### Genetic structure

PCA based on the distance of Bruvo *et al.* (2004) showed a clear divergence among cytotypes. Individuals with the same cytotype, from all populations, clustered together in three different groups (Fig. 2). Triploids clustered in an intermediate position between diploid and tetraploid cytotypes (Fig. 2). The DAPC grouped the cytotypes into ten clusters (K = 10, Fig. S1) indicating that the six sampled populations are genetically very distinct from each other (Fig. 3). These clusters, however, did not correspond to the three cytotypes. AMOVA results were consistent with DAPC indicating that most genetic variance (55 %) is associated with individuals within populations (Table 4). The single-origin model (B) has considerably higher support than the multiple origin model(ΔAICc > 10; Table 4). Global *Gst* estimate (0.233) is consistent with DAPC and AMOVA showing significant and large genetic differentiation among populations (Fig. 3). Pairwise *Gst* ranged from -0.008 to 0.245 indicating that populations PI-M / MAR-4X and ANG-M / PD-2X are less genetically divergent than the other pairwisecombinations (Fig. S2).

**Table 4.**
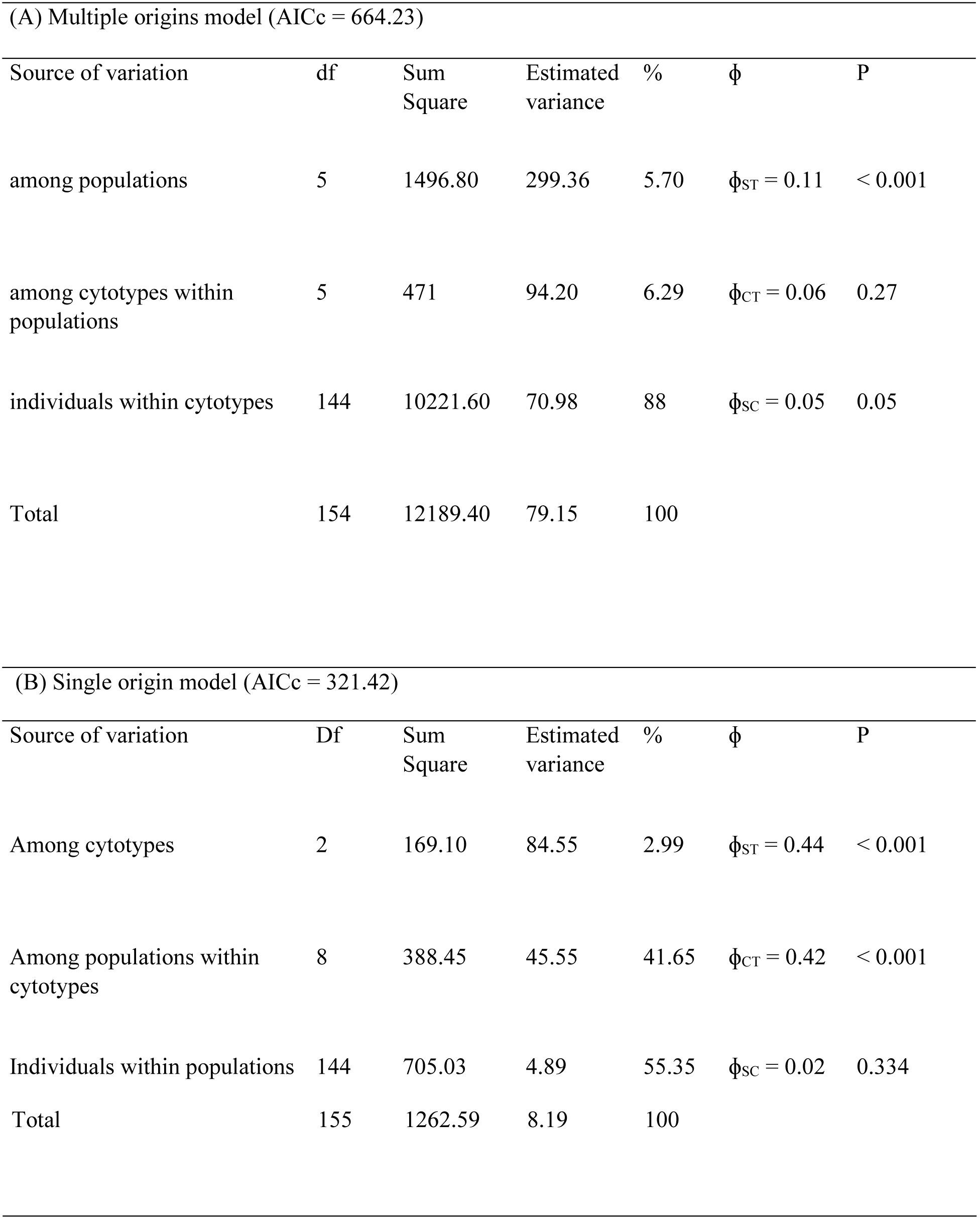
Analyses of molecular variance (AMOVA) of *Zygopetalum mackayi* Hook. populations associated with the (A) multiple origins of tetraploids and (B) single origin of polyploids. Details of AICc computation are in Table S5.

| (A) Multiple origins model (AICc = 664.23) |  |  |  |  |  |  |
| --- | --- | --- | --- | --- | --- | --- |
| Source of variation | df | Sum Square | Estimated variance | % | $\Phi$ | P |
| among populations | 5 | 1496.80 | 299.36 | 5.70 | $\Phi_{ST} = 0.11$ | < 0.001 |
| among cytotypes within populations | 5 | 471 | 94.20 | 6.29 | $\Phi_{CT} = 0.06$ | 0.27 |
| individuals within cytotypes | 144 | 10221.60 | 70.98 | 88 | $\Phi_{SC} = 0.05$ | 0.05 |
| Total | 154 | 12189.40 | 79.15 | 100 |  |  |
| (B) Single origin model (AICc = 321.42) |  |  |  |  |  |  |
| Source of variation | Df | Sum Square | Estimated variance | % | $\Phi$ | P |
| Among cytotypes | 2 | 169.10 | 84.55 | 2.99 | $\Phi_{ST} = 0.44$ | < 0.001 |
| Among populations within cytotypes | 8 | 388.45 | 45.55 | 41.65 | $\Phi_{CT} = 0.42$ | < 0.001 |
| Individuals within populations | 144 | 705.03 | 4.89 | 55.35 | $\Phi_{SC} = 0.02$ | 0.334 |

|  |  |  |  |  |
| --- | --- | --- | --- | --- |
| Total | 155 | 1262.59 | 8.19 | 100 |

**Figure 2.**
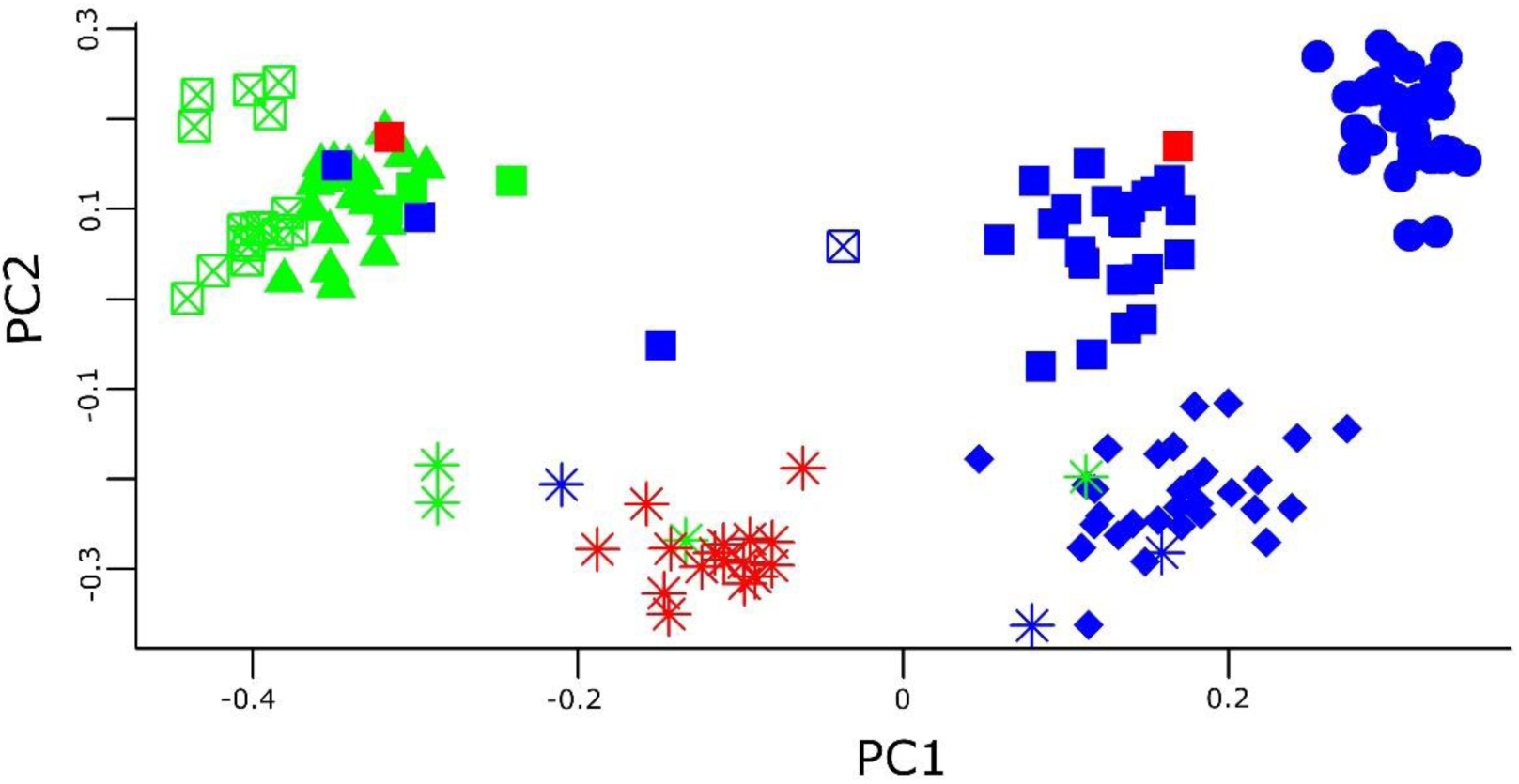
Principal component analysis of the Bruvo distance matrix of 154 MLGs of *Zygopetalum mackayi* Hook. based on eight microsatellite loci. Population codes follow Table 1: NB-2X (circles), PD-2X(diamonds), ANG-M (asterisks), PI-M (filled squares), GAR-4X (crossed squares), MAR-4X (triangles). Cytotype color follow Fig 1, diploids (blue), triploids (red), tetraploids (green). (PC1) = first principal component, (PC2) = second principal component.

**Figure 3.**
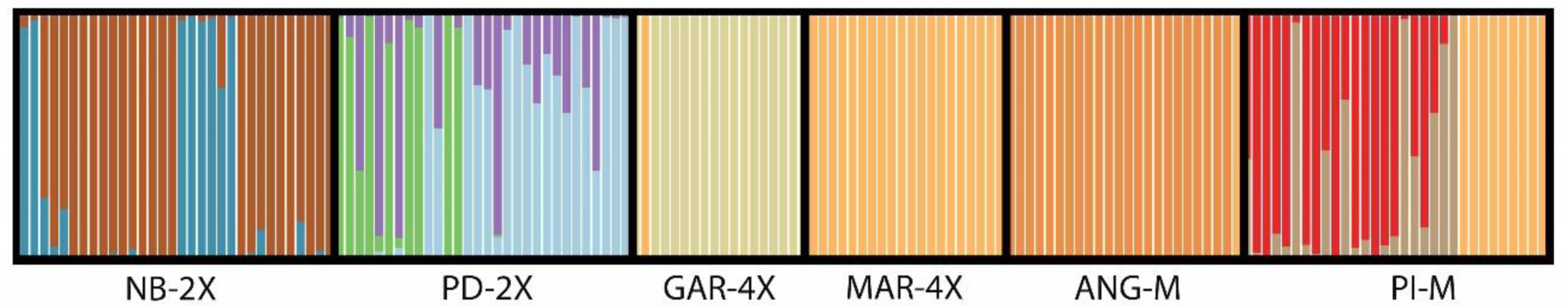
Discriminant analysis of principal components (DAPC) barplots assigning 155 accessions/cytotypes of *Zygopetalum mackayi* Hook. obtained from eight microsatellite loci based on ten inferred genetic clusters (K = 10), which are indicated by different colors. Colors in each bar represent the probability of a sampled individual to belong to a genetic cluster. Codes below bars correspond to population codes according toTable 1.

## DISCUSSION

In this study we designed microsatellite markers for individuals from six populations of the orchid *Z. mackayi* to understand the relationship between facultative apomictic reproduction and the occurrence of multiple cytotypes in this species. Specifically, we tested the hypothesis that mixed-cytotype populations of *Z. mackayi* present a higher incidence of apomictic reproduction than populations dominated by only one cytotype due to reproductive interference between sexual diploids and apomictic tetraploids. In addition, we describe the nature of contact between cytotypes in mixed populations (primary vs. secondary) and the origin of the intermediate cytotype. The results are discussed considering the role of apomixis in the evolutionary dynamics of *Z. mackayi* populations.

### The nature of cytotype contact

Our results indicate the nature of contact between diploids and tetraploids of *Z. mackayi* is secondary and that intercytotype crosses produce hybrids with intermediate chromosome number. Patterns of genetic structure and diversity clearly indicate individuals with the same chromosome number but occurring in different populations are genetically more similar to each other than to individuals with different chromosome number occurring in the same population. All triploids sampled in this study emerged as genetic intermediates between diploids and tetraploids (type I hybrids, Petit *et al.* 1999). There are no triploids present in pure diploid populations, which rules out that triploid individuals have arisen through unreduced gamete formation (Ramsey & Schemske 1998).

Our results agree with most studies available for mixed-ploidy populations which show diploids and tetraploids as dominant cytotypes resulting from secondary contact with the presence of a rare and intermediate cytotype (revised by Kolář *et al.* 2017). To our knowledge, this is the first study to confirm a secondary contact zone of mixed-cytotype species from high-elevation rocky complexes (HERCs) from Brazil. Polyploidy is expected to occur in plants from montane environments such as the Eastern Brazilian HERCs, as polyploidy maintain heterozygosity despite inbreeding (James 1992; Hopper 2009). Yet, data from other stable, nutrient-poor mountainous environments are controversial. While in the Southwest Australian Floristic Region there is a high incidence of polyploids (Hopper 1979), the Greater Cape Floristic Region, in southern Africa, has a low frequency of polyploids compared to global levels (Oberlander *et al.* 2012). Additional factors, such as the frequency of hybridization events and the high physiological cost of polyploidy in the presence of phosphorous-poor soils (Šmarda *et al.* 2013; Guinard *et al.* 2016) may be important to explain the patterns of occurrence of polyploids in these environments.

### Triploid block

Patterns of genetic diversity among populations and cytotypes of *Z. mackayi* emerged as highly structured with no evidence of relevant gene flow among populations nor introgression among cytotypes. The absence of gene flow and the high genetic diversity observed for intermediate cytotypes, despite the occurrence of facultative apomixis, suggest triploids are constantly being formed as F1 hybrids between diploids and tetraploids and that they have low fitness.

In strictly sexual, random mating species, less frequent cytotypes will be progressively removed from mixed cytotype populations because of the frequency-dependent mating disadvantage resulting from a triploid block (Marks 1966). However, if the rate of intermediate cytotype formation and intermediate cytotype fitness related to other cytotypes are high enough, the intermediate cytotype can contribute to minority cytotype establishment (Felber & Bever; Ramsey & Schemscke 1998). For example, in *Ranunculus kuepferi*, a female triploid bridge via unreduced egg cells is the major pathway toward polyploidization (Schinkel *et al.* 2017), while in *Rorippa* spp., intermediate cytotype fitness varies in different hybrid zones related to hybrid genome size (Bleeker & Matthies 2005). For *Z. mackayi*, the high frequency of intermediate cytotypes described for some populations (Gomes *et al.* 2018) added to the high genetic diversity of intermediate cytotypes reported in this study suggest that plants with intermediate cytotypes are constantly being formed. However, as we detected no evidence of clonal reproduction (almost all individuals had distinct MLGs) nor introgression between cytotypes, triploids likely have lower fitness compared to other *Z. mackayi* cytotypes. We conclude that the intermediate cytotype of *Z. mackayi* is notimportant in the establishment of minority frequency cytotypes in mixed-cytotype populations. Rather, they act as a strong reproductive barrier between progenitor cytotypes.

### The role of apomixis

We found no evidence of reproductive interference in *Z. mackayi.* Sampled populations sampled comprised of only sexual diploids (populations NB-2X and PD-2X), only facultative apomictic tetraploids (populations GAR-4X and MAR-4X) and mixed-cytotype populations (populations ANG-M and PI-M) showed high levels of genetic and genotypic diversity and, therefore, no evidence for significant asexual reproduction. Identification of residual sexuality can be difficult because similar levels of genetic variation are known to occur between apomictic polyploids and strictly sexual populations of plant species (Paun *et al.* 2006). However, several studies showed a clear genetic signature of facultative apomictic reproduction. For example, in crosses between 4*x* sexual and 6*x* apomictic *Hieracium* spp., isozyme phenotypes and chloroplast DNA haplotypes were used to identify the frequency of non-maternal offspring (Krahulcová *et al.* 2004). In contrast, the occurrence of sex and variable mechanisms of apomixis were identified by Barcaccia *et al.* (2006) for populations of *Hypericum perfloratum* based on RAPD, ISSR and AFLP fingerprints. In apomictic populations, genotypic diversity is higher among than within populations in comparison to sexual populations (Hörandl & Paun 2007). Therefore, apomictic populations are expected to have higher levels of heterozigosity and genotypic diversity (i.e. multilocus genotype diversity) within and among populations, but lower genotypic variation when compared to sexual populations (Gornall 1999, Hörandl & Paun 2007).

The analysis of molecular variance of *Z. mackayi* populations revealed that more than half of the total genetic variation (55.3%) is present within the populations, as expected for non-apomictic populations. In contrast, Dias *et al.* (2017) found 74% of genetic variation distributed among populations of obligate apomictic *Miconia albicans*. Also, the analyses of genetic diversity and estimation of the number of multilocus genotypes within all populations of *Z. mackayi* indicated similar and high levels of genetic diversity among them. Only two individuals from population GAR-4X had the same *MLG*. In contrast, Barcaccia *et al.* (2006) found multiclonal populations, lower values for Nei’s diversity index and multilocus genotypes for the facultative apomictic *Hypericum perfloratum*. Also, the analysis of codominant data for *Z. mackayi* diploids indicate it is exclusively allogamous, as expected for strictly sexual species. In sexually selfing plants or pollen-independent apomictics, uniparental reproduction allows for rapid colonization of remote areas (Baker 1965). *Z. mackayi* is pollinator dependent (Campacci *et al.* 2017) and, therefore, unable to perform self-pollination. Still, the larger geographical range of tetraploids (Gomes *et al.* 2018) suggests they are maintaining or expanding over their area of occurrence even in the presence of putatively low fitness triploids.

Geographical/ecological parthenogenesis may depend on intrinsic effects of polyploidy, rather than apomixis (Levin 1983, Alonso-Marcos *et al.* 2019). Polyploidy is hypothesized to have transformative effects in plant ecology (Ramsey & Ramsey 2014). Multiple studies in the last few years focused on polyploidy ecology and showed a significant association of polyploidy and phenological traits, plant growth pattern, and even biological interactions (Levin 1983, Ramsey & Ramsey 2014). Our results point to a strong association of ecogeographic differentiation and polyploidy, not facultative apomixis, in *Z. mackayi*. Triploids, although also apomictic in *Z. mackayi*, do not benefit from facultative apomixis because plants with odd ploidies typically have reduced fertility due to imbalanced meiotic products and the formation of aneuploid gametes (Comai 2005). Fitness of triploids may, however, vary among populations as frequency of intermediate cytotypes among populations of *Z. mackayi* is highly uneven (Gomes *et al.* 2018). Although drift may also explain uneven frequencies of cytotypes among populations, especially for montane species such as *Z. mackayi*, our results do not indicate the occurrence of inbreeding.

Plant breeding systems can have a profound effect on a species ability to occupy new areas, persist and adapt to environmental change (Richards 2003; Hojsgaard & Hörandl 2015). Recent studies have suggested sporophytic apomixis is important to plant diversification in the Brazilian savannahs (e.g. Mendes-Rodrigues *et al.* 2011; Sampaio *et al.* 2013), but for *Z. mackayi*, apomixis is likely a consequence of polyploidy. Strictly sexual diploids and facultative apomictic polyploids meet in a secondary contact zone. The occurrence of higher fitness tetraploids added to the role of triploids as a reproductive barrier may play a crucial role in the geographical expansion of apomictic tetraploids of *Z. mackayi*. Further studies on paternity analyses of intercytotype crosses and comparative fitness of *Z. mackayi* cytotypes are underway to better understand the role of apomixis and polyploidy in this species.

## Supporting information

All supplemental data is in this file

## ACKNOWLEDGMENTS

We thank T. Campacci for collecting plant specimens in the field and for helping with DNA extractions; the staff of Laboratório de Análises Genéticas e Moleculares (LAGM/UNICAMP), especially F. Alves, R. Ferreira and A. Moraes, for the technical support, and B. Leal, E. Marsola, F. Pinheiro, M.I. Zucchi for comments on previous versions of this manuscript. This study was financed in part by the Coordenação de Aperfeiçoamento de Pessoal de Nível Superior (CAPES-FAEPEX-UNICAMP-88887.373880/2019-00) and by São Paulo Research Foundation (FAPESP, grant 2014/04426-5). Collection of plant specimens and transportation were granted by IBAMA / SISBIO (#32837). AA-P thanks FAPESP (2018/00036-9) for a post-doctoral scholarship.

## REFERENCES

Afzelius K. (1959) Apomixis and polyembryony in *Zygopetalum mackayi* Hook. Acta Horti Bergiami, 19, 7–13.

Alonso-Marcos H., Nardi F. D., Scheffknecht S., Tribsch A., Hülber K., Dobeš C. (2019) Difference in reproductive mode rather than ploidy explains niche differentiation in sympatric sexual and apomictic populations of *Potentilla puberula*. Ecology and Evolution, 9, 3588–3598.

Billotte N., Lagoda P. J. L., Risterucci A. M., Baurens F. C. (1999) Microsatellite-enriched libraries: applied methodology for the development of SSR markers in tropical crops. Fruits, 54, 277–288.

Bleeker W., Matthies A. (2005) Hybrid zones between invasive *Rorippa austriaca* and native *R. sylvestris* (Brassicaceae) in Germany: ploidy levels and patterns of fitness in the field. Heredity, 94, 664 – 670.

Botstein D., White R. L., Skolnick M., Davis R. W. (1980) Construction of a genetic linkage map in man using restriction fragment length polymorphisms. American journalof human genetics, 32, 314–331.

Barcaccia G., Arzenton F., Sharbel T. F., Varotto S., Parrini P., Lucchin M. 2006. Genetic diversity and reproductive biology in ecotypes of the facultative apomict *Hypericum perforatum* L. Heredity, 96, 322–334.

Bruvo R., Michiels N. K., D’Souza T. G., Schulenburg H. (2004) A simple method for the calculation of microsatellite genotype distances irrespective of ploidy level. Molecular Ecology, 13, 2101–2106.

Burnham K. P., Anderson D. R. (1998) Practical use of the information-theoretic approach. In: Burnham K. P., Anderson D. R. (Eds), Model Selection and Inference,Springer, New York, USA: 75–117.

Campacci T. V. S., Castanho C. T., Oliveira R. L. F., Suzuki R. M., Catharino E. L. M., Koehler S. (2017) Effects of pollen origin on apomixis in *Zygopetalum mackayi*orchids. Flora-Morphology, Distribution, Functional Ecology of Plants, 226, 96–103.

Clark L. V., Schreier A. D. (2017) Resolving microsatellite genotype ambiguity in populations of allopolyploid and diploidized autopolyploid organisms using negative correlations between allelic variables. Molecular Ecology Resources, 17, 1090–1103.

Comai L. (2005) The advantages and disadvantages of being polyploid. Nature reviews genetics, 6, 836–846.

Cornuet J. M., Luikart G. (1996) Description and power analysis of two tests for detecting recent population bottlenecks from allele frequency data. Genetics, 144, 2001–2014.

Creste S., Tulmann Neto A., Figueira A. (2001) Detection of single sequence repeat polymorphisms in denaturing polyacrylamide sequencing gels by silver staining. Plant Molecular Biology Reporter, 19, 299–306.

Dias A. C. C., Serra A. C., Sampaio D. S., Borba E. L., Bonetti A. M., Oliveira P. E. (2018) Unexpectedly high genetic diversity and divergence among populations of the apomictic Neotropical tree *Miconia albicans*. Plant Biology, 20, 244–251.

Excoffier L., Smouse P. E., Quattro J. M. (1992) Analysis of molecular variance inferred from metric distances among DNA haplotypes: application to human mitochondrial DNA restriction data. Genetics, 131, 479–491.

Felber F. (1991) Establishment of a tetraploid cytotype in a diploid population: effect of relative fitness of the cytotypes. Journal of Evolutionary Biology, 4, 195–207.

Felber F., Bever J. D. (1997) Effect of triploid fitness on the coexistence of diploids and tetraploids. Biological Journal of the Linnean Society, 60, 95–106.

Gomes S. S. L., Vidal J. D., Neves C. S., Zorzatto C., Campacci T. V. S., Lima A. K., Koehler S. Viccini L. F. (2018) Genome size and climate segregation suggest distinct colonization histories of an orchid species from Neotropical high-elevation rocky complexes. Biological Journal of the Linnean Society, 124, 456–465.

Gornall R. J. (1999) Population genetic structure in agamospermous plants. Syistematics Association Special Volume, 57, 118–138.

Hersh E., Grimm J., Whitton J. (2016) Attack of the clones: reproductive interference between sexuals and asexuals in the *Crepis* agamic complex. Ecology and Evolution, 6, 6473–6483.

Hojsgaard D., Hörandl E. (2015) Apomixis as a Facilitator of Range Expansion and Diversification in Plants. In: Pontarotti P. (Eds), Evolutionary Biology: Biodiversification from Genotype to Phenotype, Springer; Berlin: 305–327

Hopper S. D. (1979) Biogeographical aspects of speciation in the southwest Australian flora. Annual Review of Ecology and Systematics, 10, 399–422.

Hopper S. D. (2009) OCBIL theory: towards an integrated understanding of the evolution, ecology and conservation of biodiversity on old, climatically buffered, infertile landscapes. Plant and Soil, 322, 49–86.

Hörandl E., Paun O. (2007) Patterns and sources of genetic diversity in apomictic plants: implications for evolutionary potentials. In: Hörandl E., Grossniklaus U., Van Huang X., Madan A. (1999) CAP3: A DNA sequence assembly program. Genome research, 9, 868–77.

Husband B. C. (2004) The role of triploid hybrids in the evolutionary dynamics of mixed-ploidy populations. Biological Journal of the Linnean Society, 82, 537–546.

Husband B.C., Baldwin S.J., Suda J. (2013) The Incidence of Polyploidy in Natural Plant Populations: *Major Patterns and Evolutionary Processes*. In: Greilhuber J., James S. H. (1992) Inbreeding, self-fertilisation, lethal genes and genomic coalescence. Heredity, 68, 449–456.

Jombart T. (2008) adegenet: a R package for the multivariate analysis of genetic markers. Bioinformatics, 24, 1403–1405.

Jombart T., Devillard S., Balloux F. (2010) Discriminant analysis of principal components: a new method for the analysis of genetically structured populations. BMC Genetics, 11, 94 – 108.

Kamvar Z. N., Tabima J. F., Grünwald N. J. (2014) Poppr: an R package for genetic analysis of populations with clonal, partially clonal, and/or sexual reproduction. PeerJ, 2, e281.

Karunarathne P., Schedler M., Martínez E. J., Honfi A. I., Novichkova A., Hojsgaard D. (2018) Intraspecific ecological niche divergence and reproductive shifts foster cytotype displacement and provide ecological opportunity to polyploids. Annals of Botany, 121, 1183–1196.

Keenan K., McGinnity P., Cross T. F., Crozier W. W., Prodöhl P. A. (2013) diveRsity: An R package for the estimation and exploration of population genetics parameters and their associated errors. Methods in Ecology and Evolution, 4, 782–788.

Kearney M. 2005. Hybridization, glaciation and geographical parthenogenesis. Trends in Ecology & Evolution, 20, 495–502.

Köhler C., Scheid O. M., Erilova A. (2010) The impact of the triploid block on the origin and evolution of polyploid plants. Trends in Genetics, 26, 142–148.

Kolář F., Fér T., Štech M., Trávníček P., Dušková E., Schönswetter P., Suda J. (2012) Bringing together evolution on serpentine and polyploidy: spatiotemporal history of the diploid-tetraploid complex of *Knautia arvensis* (Dipsacaceae). PLoS One, 7, e39988.

Kolář F., Čertner M., Suda J., Schönswetter P., Husband B. C. (2017) Mixed-ploidy species: progress and opportunities in polyploid research. Trends in Plant Science, 22, 1041–1055.

Krahulcová A., Papoušková S., Krahulec F. (2004) Reproduction mode in the allopolyploid facultatively apomictic hawkweed *Hieracium rubrum* (Asteraceae, H. subgen. Pilosella). Hereditas, 141, 19–30.

Kyogoku D. 2015. Reproductive interference: ecological and evolutionary consequences of interspecific promiscuity. Population Ecology, 57, 253–260.

Levin D. A. (1983) Polyploidy and novelty in flowering plants. The American Naturalist, 122, 1–25.

Levin D. A. (1975) Minority cytotype exclusion in local plant populations. Taxon, 24, 35–43.

Luikart G., Cornuet J. M. (1998) Empirical evaluation of a test for identifying recently bottlenecked populations from allele frequency data. Conservation biology, 12, 228–237.

Marks G. E. (1966) The enigma of triploid potatoes. Euphytica, 15, 285–290.

Markgraf V., McGlone M., Hope G. 1995. Neogene paleoenvironmental and paleoclimatic change in southern temperate ecosystems - a southern perspective. Trends in Ecology & Evolution, 10, 143–147.

Mendes-Rodrigues C., Ranal M. A., Oliveira P. E. (2011) Does polyembryony reduce seed germination and seedling development in *Eriotheca pubescens* (Malvaceae: Bombacoideae)? American Journal of Botany, 98, 1613–1622.

Nei M. (1978) Estimation of average heterozygosity and genetic distance from a small number of individuals. Genetics, 89, 583–590.

Nei M., Chesser R. K. (1983) Estimation of fixation indices and gene diversities. Annals of Human Genetics, 47, 253–259.

Oberlander K. C., Dreyer L. L., Goldblatt P., Suda J., Linder H. P. (2016) Species-rich and polyploid-poor: Insights into the evolutionary role of whole-genome duplication from the Cape flora biodiversity hotspot. American Journal of Botany, 103, 1336–1347.

Paun O., Stuessy T. F., Hörandl E. (2006) The role of hybridization, polyploidization and glaciation in the origin and evolution of the apomictic *Ranunculus cassubicus* complex. New Phytologist, 171, 223–236.

Petit C., Bretagnolle F., Felber F. (1999) Evolutionary consequences of diploid–polyploid hybrid zones in wild species. Trends in Ecology Evolution, 14, 306–311.

Piry S., Luikart G., Cornuet J. M. (1999) BOTTLENECK: a computer program for detecting recent reductions in the effective population size using allele frequency data. Journal of Heredity, 90, 502–503.

R Core Team. 2017. dR: A Language and Environment for Statistical Computing. https://www.R-project.org/

Ramsey J., Schemske D. W. (1998) Pathways, mechanisms, and rates of polyploid formation in flowering plants. Annual Review of Ecology and Systematics, 29, 467–501.

Ramsey J., Ramsey T. S. 2014. Ecological studies of polyploidy in the 100 years following its discovery. Philosophical Transactions of the Royal Society B: Biological Sciences, 369, 2013.0352.

Ribeiro J. M. C., Spielman A. 1986. The Satyr effect: a model predicting parapatry and species extinction. American Naturalist, 128, 513–528.

Richards A. J. (2003) Apomixis in flowering plants: an overview. Philosophical Transactions of the Royal Society of London. Series B: Biological Sciences, 358, 1085–1093.

Rushworth C. A., Windham M. D., Keith R. A., Mitchell-Olds T. (2018) Ecological differentiation facilitates fine-scale coexistence of sexual and asexual *Boechera*. American Journal of Botany, 105, 2051–2064.

Sampaio D. S., Bittencourt Júnior N. S., Oliveira P. E. (2013) Sporophytic apomixis in polyploid *Anemopaegma* species (Bignoniaceae) from Central Brazil. Botanical Journal of the Linnean Society, 173, 77–91.

Sampson J. F., Byrne M. (2011) Genetic diversity and multiple origins of polyploid *Atriplex nummularia* Lindl. (Chenopodiaceae). Biological Journal of the Linnean Society, 105, 218–230.

Schinkel C. C., Kirchheimer B., Dullinger S., Geelen D., De Storme N., Hörandl E. (2017) Pathways to polyploidy: indications of a female triploid bridge in the alpine species *Ranunculus kuepferi* (Ranunculaceae). Plant Systematics and Evolution, 303, 1093–1108.

Servick S., Visger C. J., Gitzendanner M. A., Soltis P. S., Soltis D. E. (2015) Population genetic variation, geographic structure, and multiple origins of autopolyploidy in *Galax urceolata*. American Journal of Botany, 102, 973–982.

Shannon C. E. (2001) A mathematical theory of communication. ACM SIGMOBILE Mobile Computing and Communications Review, 5, 3–55.

Simpson E. H. (1949) Measurement of diversity. Nature, 163, 688.

Šmarda P., Hejcman M., Březinová A., Horová L., Steigerová H., Zedek F., Bureš P., Hejcmanová P., Schellberg J. (2013) Effect of phosphorus availability on the selection of species with different ploidy levels and genome sizes in a long-term grassland fertilization experiment. New Phytologist, 200, 911–921.

Trávnícek P., Kubátová B., Čurn V., Rauchová J., Krajníková E., Jersáková J., Suda J. (2011). Remarkable coexistence of multiple cytotypes of the *Gymnadenia conopsea* aggregate (the fragrant orchid): Evidence from flow cytometry. Annals of Botany, 107, 77–87.

Temnykh S., DeClerck G., Lukashova A., Lipovich L., Cartinhour S., McCouch S. (2001) Computational and experimental analysis of microsatellites in rice (*Oryza sativa* L.): frequency, length variation, transposon associations, and genetic marker potential. Genome Research, 11, 1441–1452.

Verduijn M. H., Van Dijk P. J., Van Damme J. M. M. (2004) The role of tetraploids in the sexual–asexual cycle in dandelions (*Taraxacum*). Heredity, 93, 390–398.

Zhao S., Guo Y., Sheng Q., Shyr Y. (2014) Heatmap3: an improved heatmap package with more powerful and convenient features. BMC Bioinformatics, 15, 16.

