## Supplementary material for "Secondary origin, hybridization and sexual reproduction in a diploid- tetraploid contact zone of the facultative apomictic orchid *Zygopetalum mackayi*": All supplemental data is in this file

**Table S1**. Synopsis of characteristics of 56 nuclear SSR loci isolated from *Zygopetalum mackayi* Hook.

| Library | primer name | motif | sequence (F;R) | product size (pb) |
|---|---|---|---|---|
| NB(ZM398) | NB1 | (GT)7 | CGCGCTTACAATGCAATTAG<br>CTGACTTTTCCCGAGCCTTA | 238 |
| NB(ZM398) | NB2 | (GA)23 | TCCCTCCATATTCCCTCTCC<br>GCCAAAAAGTTTCCAAAAAGTG | 260 |
| NB(ZM398) | NB3 | (TG)6 | GAGTGAGAGTGCGTGCAT<br>CACACACACAAAACACATAGAT | 123 |
| NB(ZM398) | NB4 | (TG)6(GT)7 | TTTGGATTGTGTGTCATTTGTG<br>CTCCATCTCTCACACACCTCA | 195 |
| NB(ZM398) | NB5 | (TG)8(GA)9 | TCTTGCTTACGCGTGGACTA<br>CACTCTCACACCCACAAAGAAA | 102 |
| NB(ZM398) | NB6 | (GT)13 | CGCATGCGTGAATTAGTCTG<br>TCTTGCTTACGCGTGGACTA | 132 |
| NB(ZM398) | NB7 | (TG)5(GT)7(AG)7(TG)5 | TGTGGTTGTATGTGTATGTCCT<br>CACACACTGGTAAACACACACT | 142 |
| NB(ZM398) | NB8 | (TG)10.. (GA)10 | CTGCATGTATGTTTGTGCATGT<br>CACACACACATTCACTCCCTCT | 133 |
| NB(ZM398) | NB9 | (TG)12(GT)5 | TGTAGGTTTGGATTTGTGTA<br>ACATCCTCACTCACTCACAC | 111 |
| NB(ZM398) | NB10 | (AC)8(AG)18 | AATCCTTCCCAAGGAGCAAT<br>TCTTGCTTACGCGTGGACTA | 251 |
| PI(Pi-14) | PI1 | (GT)5...(AG)14(GT)5 | GCGTCTATGCAGCTGTGTGT<br>GGCACATGCACACTCTATTCC | 151 |
| PI(Pi-14) | PI2 | (GT)8(GT)6 | TGATGTATGTGAGACCATTTGTG<br>TCACTCACTCACATAAACATGCTC | 221 |
| PI(Pi-14) | PI3 | (TG)7...(TG)6 | TTGAGTGAGAGTGTGAGCGAAT<br>CGCATACACACAGAACACATTG | 157 |
| PI(Pi-14) | PI4 | (TG)10 | GTGAGAGCTTTTTGCGTGTG<br>CACTCATTCACTCACGCTCAC | 125 |
| PI(Pi-14) | PI5 | (CA)8 | AAGACGCACAGACACCCATA<br>GTGCGTGTTTGTGTGAGTGA | 117 |
| PI(Pi-14) | PI6 | (TG)10-TT-(AG)25 | ATCGCAAAACCCATGAAAAG<br>GTTGAGTTTGAAGGACTTCTATTTG | 136 |
| PI(Pi-14) | PI7 | TG(10)-TT-(AG)25 | TGTGTAAGTTGTGTATGTAAGAGTT<br>CATGTATGCACACACACTCAT | 133 |
| PI(Pi-14) | PI8 | (TG)6(GA)5...(TG)10 | CTGAGTGTGTTTGCGCATTT<br>CCCCCTTACACACACACACA | 148 |
| PI(Pi-14) | PI9 | (TC)15 | CGCATGTCAAGTGATGGTTT<br>CCCACCACATGAGATCAGC | 294 |
| PD 22 | PD1 | (CA)6(AC)5 | ACACCCATGCACTGACACAT | 130 |

| | | | | |
|---|---|---|---|---|
| | | | TGTGAGTTTGTGCGTAGTGTG | |
| PD 22 | PD2 | (TG)22 | CCCGATTCTGTGTAGTGTTT | 151 |
| | | | CTAACAAGGCCATCAGAAGA | |
| PD 22 | PD3 | (TG)14 | ACGGATTATGCCTGAAAATCCA | 212 |
| | | | CTCTTGCTTACGCGTGGACT | |
| PD 22 | PD4 | (AC)8 | CCACACACCCACACACTCAA | 129 |
| | | | AGTGTCAATGTGTTGTGTATGCA | |
| PD 22 | PD5 | (AC)7 | ACCTTGGGCATGAGACTAGC | 274 |
| | | | ACAGCAACATTCTCTAGCACC | |
| PD 22 | PD6 | (AC)20 | TTTCATTCGGCACCCACACA | 109 |
| | | | TGTGCGTGTATATGTTTGTTGTGA | |
| PD 22 | PD7 | (CA)6 | AGGTATGATTGATAACAACCAAG | 155 |
| | | | TTTTTAGAGCTTAGCTGACTT | |
| PD 22 | PD8 | (AC)9 | GACACAAAAACTCACACGAA | 151 |
| | | | GTGTGGGTTTATGTGAGTGT | |
| PD 22 | PD9 | (CA)11(AC)5 | CTCAAGAACACACAAATAGGC | 160 |
| | | | TTACGCGTGGACTAGAT | |
| PD 22 | PD10 | (AC)7 | AAAGACATACACACAAACGC | 154 |
| | | | ACGTGTGTTGTTTGTGTTTG | |
| Marins(ZM106) | MA1 | (CA)6 | CTAGACTCACCACGCATGCA | 228 |
| | | | ACATTTGCCCTATCTGCCCC | |
| Marins(ZM106) | MA2 | (TG)6(TATG)5 | TATGTGTGCTTCTAGTGCTC | 146 |
| | | | ACTCAAATGACAGACACACT | |
| Marins(ZM106) | MA3 | (AT)5(GT)16 | TGAAGCAGACGTTTCCAGAAAC | 170 |
| | | | ACACATATACACACAAGCACAG | |
| Marins(ZM106) | MA4 | (TG)12 | GTAGTGGGATTAAGGCTTGT | 260 |
| | | | ATCACCTGGGTGGAATTTTT | |
| Marins(ZM106) | MA5 | (AC)14 | CCACACATACTCACTCACTC | 133 |
| | | | GAGAGTGCATGTCTGTGTAT | |
| Marins(ZM106) | MA6 | (ACC)4 | GATGAAGAGGAGGAGAAACC | 258 |
| | | | CATTCCTCACTAAGGGCTAC | |
| ZM150(GAR) | GAR1 | (TG)15 | ACGCGCCTAGAAAATCCTAA | 270 |
| | | | ACCCGAAAGATCCTAGTAGG | |
| ZM150(GAR) | GAR2 | (AC)5 | GCACACACATGGACATACAA | 112 |
| | | | GCATGTTTTTGGGTGAGTGT | |
| ZM150(GAR) | GAR3 | (GT)24 | CATGCATGAGTGTGTGTGTA | 142 |
| | | | CAGGCACTCACAAACACAAA | |
| ZM150(GAR) | GAR4 | (CA)12 | GAGGATTTAATACCTACATCCT | 214 |
| | | | AAAAATCCAAAACTTCACCC | |
| ZM150(GAR) | GAR5 | (GC)9(CA)7 | ACCCACACAAACCTACTCAC | 216 |
| | | | GGGTGTGTGTGTCTGTCAAT | |
| ZM150(GAR) | GAR6 | (TG)6 | AGTGTTTTGTGAGTGGAT | 111 |
| | | | TCACAAACTTATACACAACAC | |

| | | | | |
|---|---|---|---|---|
| ZM150(GAR) | GAR7 | (AT)10(TG)8 | CAAAAAGGTGAGCTGCGTTA<br>TCTTTGGGTTCTATCCTCGC | 152 |
| ZM150(GAR) | GAR8 | (AC)9 | GCTTAGGCTCTATAAGAAAATTG<br>ACTAACAACACAACAACAGA | 306 |
| ZM150(GAR) | GAR9 | (CT)5(CA)6 | GAGACAAATCTACACACGCG<br>TGTGTGTGTGTGTCTCTCTC | 195 |
| ZM25(GAR) | GARD1 | (GT)10 | TGTGTGTGTGCATGCATTTT<br>CATAGACAAACACTCGCACA | 209 |
| ZM25(GAR) | GARD2 | (GT)5 | TTGAGTGTGTTTGTGTGAAC<br>CACACAGCTCACTAACACAT | 112 |
| ZM25(GAR) | GARD3 | (TG)25(AG)(GT)8 | TCATCCTATGTGATTTGGCAC<br>GGAGAGAATGGTTTGTAGTTCT | 190 |
| ZM25(GAR) | GARD4 | (CA)8 | ACCCCCTTGAAAATTTTGGC<br>TGTTAAGTGGTGGGCTCAAT | 129 |
| ZM25(GAR) | GARD5 | (TG)8 | TGTCCAATATGATTGAGATCA<br>TCTCAGTTTTGATGATC | 339 |
| ZM25(GAR) | GARD6 | (AC)21 | TACACGCACACAACACATAG<br>TGTATGAGTGTATGTGTGGA | 112 |
| ZM25(GAR) | GARD7 | (CA)7...(CA)5 | CTAACACACACGCACACTTG<br>GCGTGTCAATGTGTTGTGTA | 107 |
| ZM25(GAR) | GARD8 | (AC)20 | ATACACAGATGCACACAC<br>AACAGTTTTAGCTCCATACA | 261 |
| ZM25(GAR) | GARD9 | (GT)5 | TGTGTGTGTGTGTGTCTTCA<br>TTTTTGGTTGATGCGCAGG | 143 |
| ZM25(GAR) | GARD10 | (TA)6(AC)8 | AATGGGTGCTTACGCTTTTT<br>ACAGCTAACCCTGCTCTTTT | 219 |
| ZM25(GAR) | GARD11 | (TG)5 | GTATTTTTGTGTGGGTGTC<br>ACACATACCAACTACACAC | 160 |
| ZM25(GAR) | GARD12 | (TG)15 | AATCTGATGCACATTAATAATTT<br>CTAAGGCTTGTATCTCAGTT | 305 |

**Table S2.** Synopsis of characteristics of eight nuclear SSR loci isolated from *Zygopetalum mackayi* Hook. and used in this study. PIC = polymorphism information content.

| primer name | geneBank ID | motif | sequence (F;R) | product Size (pb) | PIC |
|---|---|---|---|---|---|

| | | | | | |
|---|---|---|---|---|---|
| NB2 | MK753245 | (GA)23 | TCCCTCCATATTCCCTCTCC<br>GCCAAAAAGTTTCCAAAAAGTG | 260 | 0,79 |
| NB8 | MK753246 | (TG)10.. (GA)10 | CTGCATGTATGTTTGTGCATGT<br>CACACACACATTCACTCCCTCT | 133 | 0,61 |
| PI1 | MK753247 | (GT)5...(AG)14(GT)5 | GCGTCTATGCAGCTGTGTGT<br>GGCACATGCACACTCTATTCC | 151 | 0,69 |
| PI3 | MK753248 | (TG)7...(TG)6 | TTGAGTGAGAGTGTGAGCGAAT<br>CGCATACACACAGAACACATTG | 157 | 0,84 |
| PI9 | MK753249 | (TC)15 | CGCATGTCAAGTGATGGTTT<br>CCCACCACATGAGATCAGC | 294 | 0,77 |
| MA6 | MK753250 | (ACC)4 | GATGAAGAGGAGGAGAAACC<br>CATTCCTCACTAAGGGCTAC | 258 | 0,7 |
| GAR1 | MK753251 | (TG)15 | ACGCGCCTAGAAAATCCTAA<br>ACCCGAAAGATCCTAGTAGG | 270 | 0,86 |
| GARD2 | MK753252 | (GT)5 | TTGAGTGTGTTTGTGTGAAC<br>CACACAGCTCACTAACACAT | 112 | 0,53 |

**Table S3.** Number of alleles per locus observed in each population of *Zygopetalum mackayi* Hook. sampled.

| Primer/population code | NB-2X | PD-2X | ANG-M | GAR-4X | MAR-4X | PI-M | total |
|---|---|---|---|---|---|---|---|
| NB2 | 3 | 9 | 7 | 4 | 7 | 7 | 37 |
| MA6 | 3 | 4 | 3 | 7 | 5 | 5 | 27 |
| PI9 | 2 | 4 | 6 | 3 | 5 | 8 | 28 |
| GAR1 | 2 | 8 | 7 | 10 | 5 | 6 | 38 |
| GARD2 | 2 | 2 | 2 | 4 | 2 | 4 | 16 |
| NB8 | 3 | 7 | 4 | 4 | 2 | 6 | 26 |

| | | | | | | | |
|---|---|---|---|---|---|---|---|
| PI1 | 9 | 8 | 5 | 6 | 7 | 5 | 40 |
| PI3 | 6 | 2 | 4 | 4 | 3 | 6 | 25 |
| total | 30 | 44 | 38 | 42 | 36 | 47 | 237 |

**Table S4.** Results of bottleneck tests for the diploid populations NB-2X and PD-2X and diploid individuals of population PI-M of *Zygopetalum mackayi* Hook. considering a stepwise mutation model (SMM) and infinite alleles model (IAM). Expected heterozygosity in equilibrium mutation-drift (Heq); (*+) = p < 0,05.

| population | NB-2X | | | |
|---|---|---|---|---|
| | I.A.M | | SMM | |
| locus | Heq | p value | Heq | p value |
| NB2 | 0.354 | 0.4320 | 0.476 | 0.1340 |
| PI9 | 0.355 | 0.2050 | 0.479 | 0.0270* |
| MA6 | 0.203 | 0.0220* | 0.248 | 0.0240* |
| GA1 | 0.216 | 0.3460 | 0.247 | 0.4090 |
| GA2 | 0.204 | 0.1270 | 0.246 | 0.1670 |
| NB8 | 0.346 | 0.2010 | 0.471 | 0.4540 |
| PI1 | 0.745 | 0.3760 | 0.837 | 0.0160* |
| PI3 | 0.626 | 0.0120* | 0.745 | 0.0830 |
| all loci | | 0.318 | | 0.4716 |

| population | PD-2X | | | |
|---|---|---|---|---|
| | I.A.M | | SMM | |
| locus | Heq | p value | Heq | p value |
| NB2 | 0.753 | 0.5050 | 0.838 | 0.0630 |
| PI9 | 0.486 | 0.2560 | 0.610 | 0.0500 |
| MA6 | 0.478 | 0.1480 | 0.614 | 0.4640 |
| GA1 | 0.734 | 0.0330* | 0.822 | 0.0000* |
| GA2 | 0.209 | 0.0460 | 0.257 | 0.0700 |
| NB8 | 0.708 | 0.3730 | 0.795 | 0.0490* |
| PI1 | 0.774 | 0.3180 | 0.855 | 0.0110* |
| PI3 | 0.212 | 0.0990 | 0.242 | 0.1330 |
| all loci | | 0.5183 | | 0.2386 |

| population | PI-M | | | |
|---|---|---|---|---|
| | I.A.M | | SMM | |
| locus | Heq | p value | Heq | p value |
| NB2 | 0.695 | 0.2740 | 0.792 | 0.0210* |
| PI9 | 0.211 | 0.5360 | 0.253 | 0.3940 |
| MA6 | 0.749 | 0.2870 | 0.826 | 0.2060 |
| GA1 | 0.578 | 0.4220 | 0.696 | 0.0770 |
| GA2 | 0.361 | 0.1740 | 0.487 | 0.3720 |
| NB8 | 0.636 | 0.2860 | 0.751 | 0.2500 |
| PI1 | 0.689 | 0.1480 | 0.788 | 0.3930 |
| PI3 | 0.384 | 0.1020 | 0.492 | 0.2500 |

| | | |
|---|---|---|
| all loci | 0.4949 | 0.0627 |

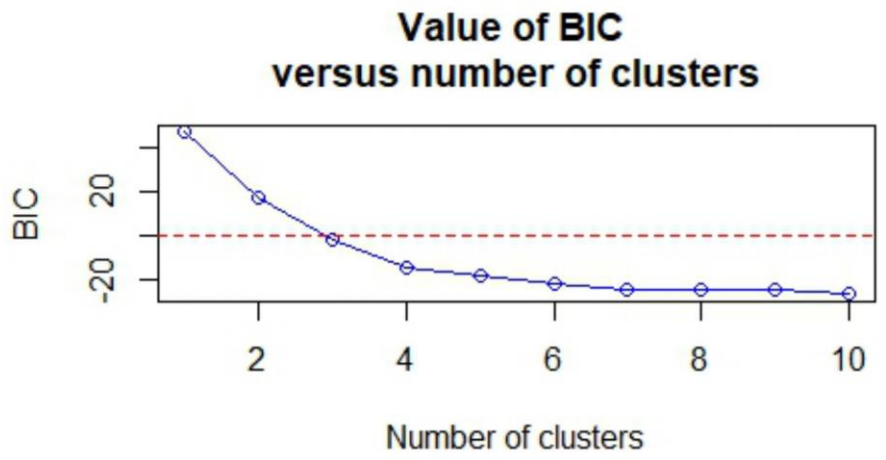


**Figure S6.** Result of K-means clustering algorithm indicating 10 as the optimal number of groups maximizing the variation between groups of *Zygopetalum mackayi* Hook. This optimal number of group was used in the DAPC to define the genetic clusters.

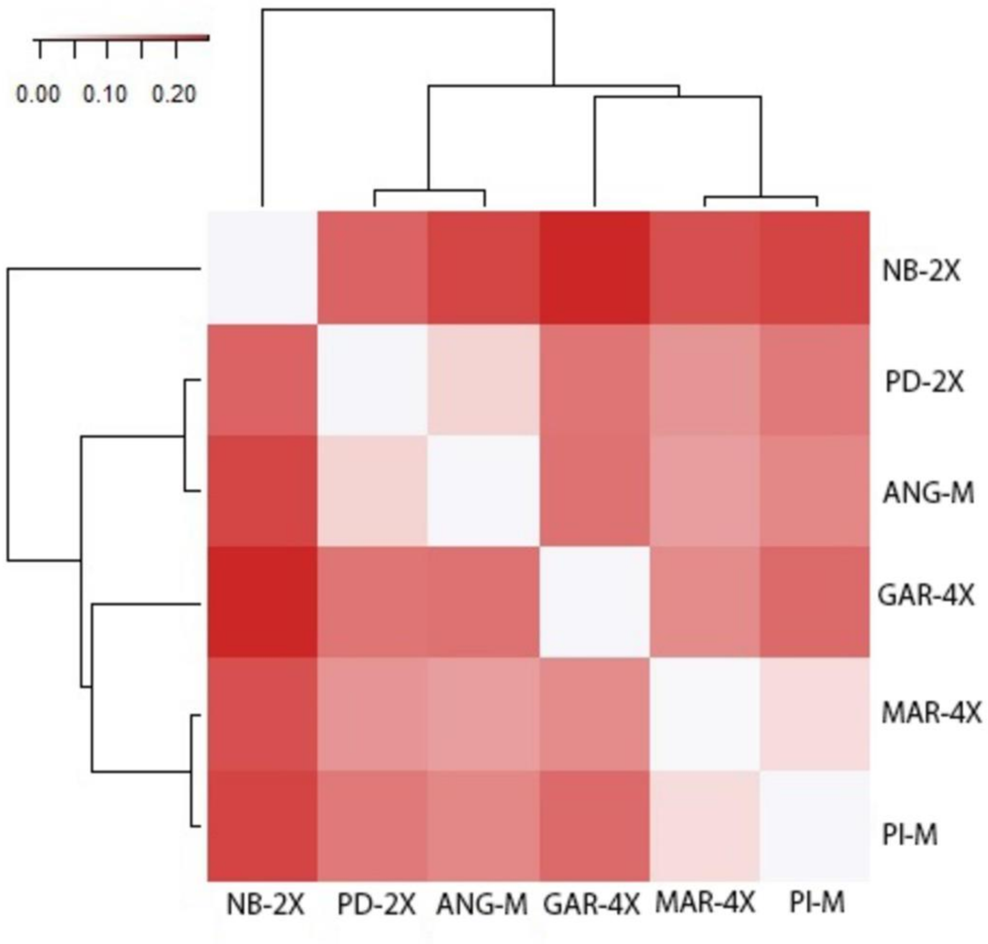


**Figure S7.** Heatmap of Gst pairwise values for populations of *Zygopetalum mackayi* Hook. sampled in this study. Population codes follow Table 1.
